# The impact of social complexity on the efficacy of natural selection in termites

**DOI:** 10.1101/2024.04.26.591327

**Authors:** Camille Roux, Alice Ha, Arthur Weyna, Morgan Lodé, Jonathan Romiguier

## Abstract

In eusocial species, reproduction is monopolized by a few reproductive individuals. From the perspective of population genetics, this implies that the effective population size (*Ne*) of these organisms is likely to be smaller compared to solitary species, as has been proposed in the literature for eusocial hymenoptera. In this study, we investigate the genomic consequences of eusociality in termites (Isoptera) on two different timescales. First, by analyzing transcriptome data from 66 Blattodea species, we focus on the ratio of non-synonymous to synonymous mutations (*d_N_/d_S_*) as a marker of natural selection efficiency and effective population size. Our results demonstrate an elevated *d_N_/d_S_* ratio in termites compared to other members of Blattodea, further generalizing the idea that convergent evolution toward eusociality strongly reduces the effective population size and the genome-wide efficiency of natural selection. Then, by comparing 68 termite transcriptomes, we show that this decrease in natural selection efficiency is even more pronounced in termites displaying high levels of social complexity. This study contributes to understanding the complex interplay between social structures and natural selection patterns, highlighting the genetic footprint of eusociality in shaping the evolution of termites.

## Introduction

Eusociality represents the highest level of social organization within the animal kingdom, characterized by a clear division of reproductive labor among different castes, cooperative care for young and the presence of overlapping generations within a colony (Keller and Chapuisat, 2001). Various forms of social organizations have emerged independently in several taxa, such as Coleoptera, Thysanoptera (Choe and Crespi, 1997), crustaceans (Duffy, 1996), spiders (Lubin and Bilde, 2007), or mammals (Jarvis et al., 1994), but the most complex forms of eusociality appeared only within Hymenoptera (e.g. bees, wasps, ants) and Isoptera (termites; Howard and Thorne, 2011). In social insects such as ants and termites, a caste is defined as a subset of individuals within a colony that perform specialized roles, often with distinct morphological traits. Both ants and termites display sharp contrasts in terms of levels of social complexity, ranging from small colonies of hundreds of individuals with low caste polymorphism to large colonies of several millions of individuals and extreme caste differentiations.

For a given number of individuals that make up a population, the shift from solitary living to eusociality corresponds to a notable decrease in the effective population size *Ne*, mainly because reproduction becomes limited to a reduced number of individuals within a colony (Romiguier, Lourenco, et al., 2014). Such reduction of *Ne* has important consequences for genome evolution due to the interaction between natural selection efficiency and random genetic drift (Kimura, 1983). The potency of selection to promote the fixation of advantageous mutations or to remove deleterious ones depends on the selection coefficient *vis-à-vis* the influence of genetic drift, whose intensity is increased for lower *N_e_* values. When the influence of selection weakens in comparison to genetic drift (*|N_e_.s| <* 1, *s* being the selective coefficient), alleles behave as if they are effectively neutral. Thus, genomes of species with a large effective population size are characterized by a higher relative accumulation rate of slightly advantageous mutations compared to species with a small *N_e_*(Galtier, 2016). In the same direction, weakly deleterious mutations are not effectively eliminated from the genomes of low *N_e_*species because genetic drift is stronger than purifying selection for them (Galtier, 2016).

This phenomenon sets a theoretical threshold dubbed the “drift barrier” (Lynch and Walsh, 2007), delineating the boundary where random genetic drift limits the ability of selection to forestall the fixation of sub-optimal alleles (Kimura et al., 1963; Ohta, 1973). The empirical demonstration of the drift barrier has established it as a key factor shaping the genetic architectures of living organisms. The central involvement of the *N_e_* effect has since been empirically demonstrated to explain the evolution of different genomic components: mutation rate (Lynch, Ackerman, et al., 2016), intron emergence (Lynch, 2002), precision of intron splicing (Benitiere et al., 2022), optimization of codon usage (Galtier et al., 2018), complexity of protein-protein interactions (Lynch, 2012), protein optimization (Huber et al., 2017) and rate of adaptive evolution (Galtier, 2016).

The evolution of coding regions themselves is shaped by the mutation-selection-drift equilibrium, with life history traits significantly impacting the rates of nonsynonymous substitutions relative to synonymous substitutions (*d_N_/d_S_*; Botero-Castro et al., 2017; Figuet et al., 2016; Nikolaev et al., 2007; Popadin et al., 2007; Rolland et al., 2020; Romiguier, Ranwez, et al., 2013; Weyna and Romiguier, 2021). At synonymous positions, the majority of mutations are presumed to be neutral, although some exceptions exist (Galtier et al., 2018). This implies that the substitution rate *d_S_* is a result of the number of mutations emerging per generation in the population (2*.N_e_.µ*, where *µ* represents the mutation rate per nucleotide per generation) multiplied by the probability of fixing a new neutral mutation 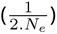. Hence, in absence of selective pressure on synonymous positions, the evolution of *d_S_*is predominantly driven by the mutation rate. Conversely, the fixation probability of new nonsynonymous mutations is directly reliant on the *N_e_.s* product: the smaller the *N_e_*, the greater the accumulation of weakly deleterious nonsynonymous mutations in genomes. Consequently, any life-history trait impacting *N_e_*is likely to increase the relative amount of deleterious nonsynonymous mutations within genomes by intensifying the role of genetic drift

Eusociality in ants, bees, wasps, termites, shrimps and spiders appears to be such a life-history trait by restricting reproduction to few individuals. In spiders of the genus *Stegodyphus*, where only a small fraction of females lay eggs, whole-genome analyses have shown that the shift towards higher sociality is linked to a continuous reduction in effective population size and an increase in *d_N_/d_S_* (Ma et al., 2024). However, the analysis of 169 Hymenopteran species provide a slightly nuanced view of the link between eusociality and effective population size (Weyna and Romiguier, 2021). While social Hymenoptera featured an increase of *d_N_/d_S_* compared to solitary species, the results highlighted the notable exception of solitary bees displaying the same level of *d_N_/d_S_* than their social counterparts. Also, another recent study suggests that low division of reproductive labour (i.e. worker reproduction) may not necessarily translate into lower *d_N_/d_S_* (i.e increased strength of natural selection, Barkdull and Moreau (2023)). These results highlighted that strong decreases of effective population size is not necessarily a direct consequence of division of reproductive labour, but might as well predate it.

In this study, we aim to ascertain whether the negative impact of eusociality on the efficacy of natural selection applies to termites. Similarly to ants despite *≈* 383 my of divergence (Kumar et al., 2017), termites features various level of eusociality, the most complex forms being characterized by particularly large colonies (up to *>* 10^6^ individuals; Porter and Hawkins, 2001) and high caste specialization. We thus hypothesize further that termite species with the highest levels of social organization might feature the lowest effective population size. This would mean that large superorganisms (i.e. large colonies) are analogous in terms of *Ne* with large organisms, reflecting previous works in mammals or birds where species with large body size feature high dN/dS ratio (Botero-Castro et al., 2017; Romiguier, Ranwez, et al., 2013).

To explore this inquiry, we leverage two publicly accessible sequencing datasets delineating phylogenetic information at two different evolutionary levels:

1. across Blattodea, allowing for the comparison between 6 species of termites and 60 non-termites species (Evangelista et al., 2019).
2. across 68 species of termites, to study the effect of variation in levels of social complexity on *d_N_/d_S_* (Bucek et al., 2019).

Our findings reveal that termites (*i.e.* eusocial Blattodea) exhibit higher *d_N_/d_S_* ratios than solitary Blattodea species, interpreted as a relaxation of purifying selection. This finding aligns with a recently published study that compared three termite species with five cockroach species (Ewart et al., 2024). Furthermore, among termites, species displaying the highest levels of social complexity tend to exhibit higher *d_N_/d_S_* ratio in comparison to those considered to exhibit simpler social organization.

## Material and methods

The analysis presented here focuses on two phylogenetic scales: within the Blattodea in first, then a zoom on the Isoptera. For both scales, all the data including molecular alignments, phylogenetic trees and life history traits were taken from the literature. *d_N_/d_S_* by branch was then inferred in this study, based on the available aligned transcriptomes and the reconstructed phylogenetic trees.

### Blattodea dataset

The molecular data (sequence alignments + phylogenetic tree) for the Blattodea come entirely from the study by Evangelista et al. (2019). This dataset corresponds to 3,244 coding genes aligned in 66 species, and is from the Genbank Umbrella Bioproject (ID PRJNA183205). The 66 Blattodea species had been chosen by the authors to represent the major lineages (except the Anaplectidae). Evangelista et al. (2019) sequenced the transcriptomes of 31 Blattodea species, and complemented this dataset by transcriptomic data coming from Misof et al. (2014) and Wipfler et al. (2019), consisting of 14 additional Blattodea and 21 outgroups. RNA extraction, library construction and paired-end sequencing (Illumina HiSeq 2000) are part of the 1KITE project (www. 1kite.org). The used phylogenetic tree corresponds to the file Blattodea_FigTree_95_aa_run1_clean.treefile (Supplementary Material), was produced by Evangelista et al. (2019) using a maximum-likelihood approach implemented in IQ-TREE (Haeseler et al., 2014; Kalyaanamoorthy et al., 2017). The final dataset we used for the *d_N_/d_S_* analysis is composed over the 66 surveyed species by a median number of 2,102 coding genes per species, ranging from 1,243 in *Nyctibora* to 2,573 in *Zootermopsis nevadensis*.

Before carrying out an ANOVA analysis to test an effect of social organization on *d_N_/d_S_*, we checked whether this ratio followed a normal distribution in each social group using a Shapiro-Wilk test (*P* -values ranging from 0.077 in the gregarious group to 0.838 in the eusocial group). Then we checked that variances of the *d_N_/d_S_* ratios are homogeneous between the different social systems using a Levene’s test (*P* = 0.3). Once the normality of the *d_N_/d_S_* and the homogeneity of the variances between groups had been confirmed, we performed an ANOVA test to check an effect of sociality. To identify specific differences between groups, we used a Tukey-HSD (Honest Significant Difference) test, which tests the differences in means between each pair of groups

### Isoptera dataset

The Isoptera dataset comes from the study carried out by Bucek et al. (2019). They sequenced the transcriptomes of 53 termite species and two entire genomes. These data were used to complete publicly available genomes and transcriptomes for 13 termite species and 7 dictyopteran outgroups (*Blaberus atropos*, *Blattella germanica*, *Cryptocercus wrighti*, *Empusa pennata*, *Mantis religiosa*, *Metallyticus splendidus*, *Periplaneta americana*; Bourguignon, Šobotník, et al., 2016; Dedeine et al., 2015; Harrison et al., 2018; Hayashi et al., 2013; Huang et al., 2012; Misof et al., 2014; Mitaka et al., 2016; Poulsen et al., 2014; Terrapon et al., 2014; Wu et al., 2016). Transcriptomes of workers were sequenced by Bucek et al. (2019) using the Illumina HiSeq 4000 platform. The authors then aligned the assembled orthologous genes using MAFFTv7.305 (Katoh and Standley, 2013). They inferred the phylogenetic tree using IQ-TREE 1.6.7 (Nguyen et al., 2015). The used phylogenetic tree corresponds to the file Isoptera_Bucek.treefile (Supplementary Material). The final dataset we used to estimate *d_N_/d_S_* corresponds to a median number of 3,604 coding genes per species, ranging from 2,391 (*Reticulitermes grassei*) to 3,927 (*Blattella germanica*).

### Phylogenetic estimation of the *d_N_/d_S_* ratio

*d_N_/d_S_* values were computed using the mapnh program within the TestNH suite (J Dutheil and Boussau, 2008; JY Dutheil et al., 2012; Guéguen and Duret, 2018; Romiguier, Figuet, et al., 2012; https://github.com/ BioPP/testnh) from the publicly available cleaned alignments, and from the tree topologies available for both the Blattodea and Isoptera datasets. Each locus which had sequences with over 90% non-informative sites were removed. mapnh estimates *d_N_* and *d_S_* for four substitution categories (*W → S*, *S → W* , *W → W* , *S → S*, referring to nucleotides affected by substitutions, AT = W for Weak and CG = S for Strong). Non-synonymous and synonymous substitution counts are estimated for each branch in the phylogeny. However, for downstream analyses, we use the *d_N_/d_S_* ratio estimated from the terminal branches as representative for each species. Alignments with fewer than 30 species were discarded.

### Testing for relaxation of natural selection

To assess the relaxation of selective pressure, we employed the RELAX model within the HyPhy v.2.5 software (Schrader et al., 2021; Wertheim et al., 2015). The RELAX model estimates the selection intensity parameter (*k*), allowing us to compare a null model (*k* = 1) with an alternative model, thereby evaluating the strength of natural selection.

A *k* value greater than 1 indicates intensified natural selection, while a *k* value less than 1 suggests relaxed natural selection in the test branches (six termite species) relative to the reference branches (60 non-termite species) using the dataset from Evangelista et al. (2019). We assessed the statistical significance of the *k* value (p<0.05) using a Bonferroni correction.

### Social traits

The data from Evangelista et al. (2019) were utilized to investigate potential variations in *d_N_/d_S_* within Blattodea associated with eusociality. Each species was correlated with its social category following Michener’s classification in 1969 Michener (1969) and average individual size. Sociality data were compiled from various bibliographic (Bohn, 1989; Cabrera and Scheffrahn, 2001; Legendre, 2007; Lihoreau, 2009; Picker, 2012; Roth, 1986, 2000) and online (Bugguide; Cockroachcare; Collections.museumvictoria; Jasa; Ozanimals; Padil; Roachcrossing; Virginiacheeseman; Wikipedia) sources. When species-level information was lacking, we utilized the average observation from within genera or subfamilies.

The data sourced from Bucek et al. (2019) were employed to identify potential variations in *d_N_/d_S_* within Isoptera linked to varying degrees of sociality. Each species was detailed with various characteristics, including category (Neoisoptera which is a clade grouping most families with high level of sociality vs Basal Isoptera), taxonomic family, queen(s) egg-laying count, approximate colony size, colony type (monogynous, polygynous, or both), polygyny type, worker type (“true workers” than are completely sterile vs “pseudergates” that can become reproductives), dietary habits, geographical distribution, soldier and queen morphometrics, and invasive status. Notably, the Termitidae and Rhinotermitidae families, encompassing several paraphyletic clades in the dataset, were arbitrarily divided into Termitidae A and B and Rhinotermitidae A to E to study monophyletic groups. However, this division doesn’t align with established subfamilies in taxonomy. Data on species characteristics were collected from various literature sources (Ahmad, 1963; Atkinson and ES Adams, 1997; Barbosa and Constantino, 2017; Bourguignon, Scheffrahn, et al., 2010; Chhotani, 1975; Constantino and De Souza, 1997; Constantino, 1990; De Saeger, 1954; Dejean and Fénéron, 1996; Dupont et al., 2009; Emerson and Banks, 1965; Evans, 2011; Florian et al., 2019; Fougeyrollas et al., 2015; F Gay, 1974; F Gay and Barrett, 1983; FJ Gay and JA Watson, 1982; Ghesini and Marini, 2013; Goodisman and Crozier, 2002; Hanus et al., 2006; Hellemans et al., 2017; Husseneder et al., 1998; Krishna, 1963; Krishna and CL Adams, 1982; Krishna and Araujo, 1968; Lewis, 2009; Liang et al., 2017; Maistrello and Sbrenna, 1996; Martius and Ribeiro, 1996; Matsuura, 2002; Miller, 1994; Miura et al., 2003; Noirot, 1959; Noirot, 1989; Parmentier and Roisin, 2003; Rasib and Saeed Akhtar, 2012; Scheffrahn and Krecek, 1999; Scheffrahn, Su, et al., 1999; Snyder, 1924; Soleymaninejadian et al., 2014; Thorne, 1984; Vargo, 2019; Vargo et al., 2006; J Watson, Brown, et al., 1989; J Watson, Metcalf, et al., 1977; Williams, 1966). Similar to the dataset in Evangelista et al. (2019), the search scope was extended to include closely related genera and subfamilies because of shared characteristics among closely related species.

## Results and discussion

### *d_N_/d_S_* evolution in Blattodea

We estimated the *d_N_/d_S_* ratio as a proxy of natural selection efficiency across a phylogenetic tree of 66 Blattodea species, including 6 termites (Isoptera). The dataset and phylogenetic tree employed in this investigation are derived from the study by Evangelista et al. (2019) encompassing 3,244 coding genes (Fig. 1).

**Figure 1.**
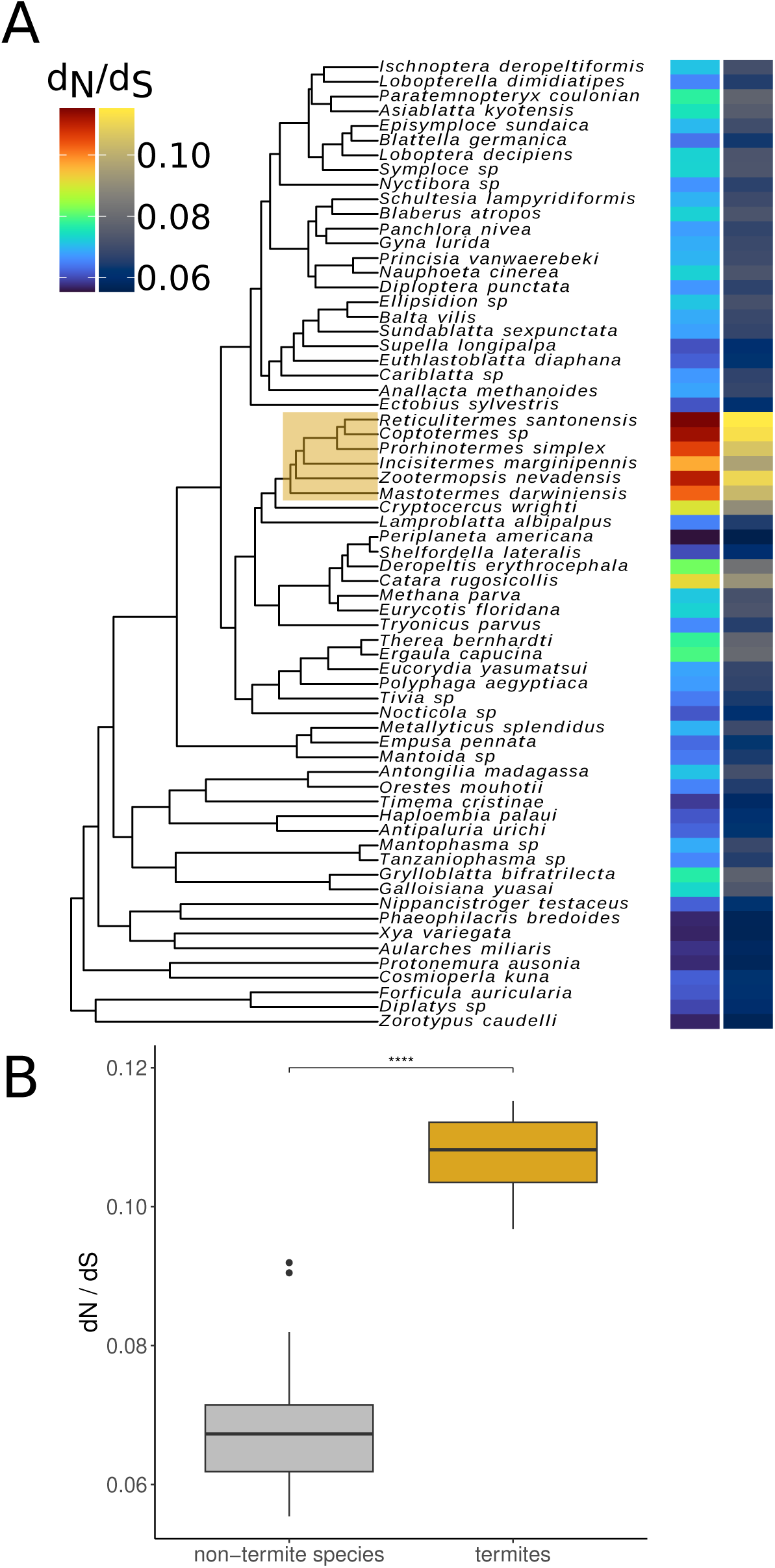
Distribution of the *d_N_/d_S_* ratio in Blattodea. A. The phylogenetic tree employed in this study was produced by Evangelista et al. (2019), constructed from sequenced transcriptomes. Utilizing the same set of coding genes, we estimated the *d_N_/d_S_* ratios along each phylogenetic branch by mapnh. In this study, we exclusively report the *d_N_/d_S_* values corresponding to the terminal branches, represented here by a gradient of colours between 0.055 and 0.115. Two color palettes are used here to include the diversity of color perception. The clade encompassing termites is highlighted by a yellow rectangle. B. Boxplot representing the distribution of *d_N_/d_S_* ratios in the terminal branches of Blattodea. In grey: estimated values for non-termite species. In yellow: estimated values for termites. Horizontal lines indicate median values. Statistical significance is denoted as follows: *ns* (*P >* 0.05), * (*P ≤* 0.05), ** (*P ≤* 0.01), *** (*P ≤* 0.001), **** (*P ≤* 0.0001).

Upon comparing the mean genomic *d_N_/d_S_* ratios calculated for the external branches leading to the 66 species under study, we observe a notably higher *d_N_/d_S_* ratio that can be interpreted as lower efficiency of natural selection within the clade that includes termites (mean=0.107; standard deviation=0.00687) as opposed to outside this group (mean=0.0675; standard deviation=0.00756). This disparity is statistically significant, as confirmed by a Wilcoxon signed-sum test (*W* = 0; *P* = 6.235 *×* 10*^−^*^5^). The two distributions do not overlap, with the highest *d_N_/d_S_* for non-termites (0.092 in *Catara rugosicollis*) being lower than the lowest *d_N_/d_S_* for termites (0.097 in *Incisitermes marginipennis*).

In mammals, body size is known to strongly influence the *d_N_/d_S_* ratio because of its negative relationship with *Ne*. Thus, large and long-lived species display on average a higher *d_N_/d_S_* ratio than small and short-lived mammals (Nikolaev et al., 2007). Here we test whether a similar relationship is found in Blattodea (Fig. 2-A; Table S1).

**Figure 2.**
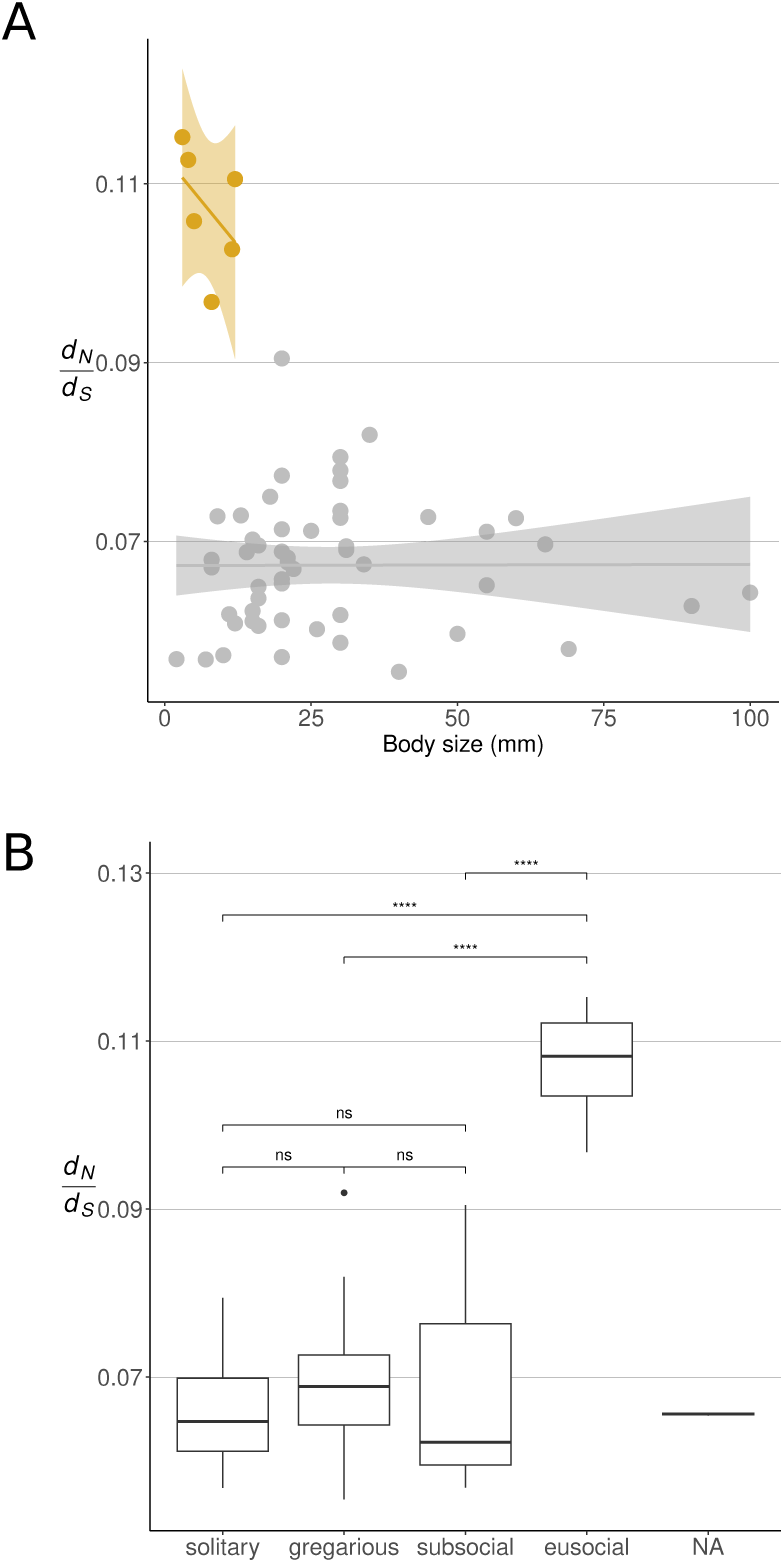
Comparative analysis of body size and social system influences on *d_N_/d_S_* ratios Panel A: relationship between organism body size reported for adults (in mm) and *d_N_/d_S_*. Termite data points are shown in orange, while non-termite points are displayed in grey. Panel B: distribution of *d_N_/d_S_* ratios within different social systems according to Michener (1969). Bonferroni-adjusted statistical significance is denoted as follows: *ns* (*P >* 0.05), * (*P ≤* 0.05), ** (*P ≤* 0.01), *** (*P ≤* 0.001), **** (*P ≤* 0.0001).

To do this, we take advantage of the information available on adult body length in Blattodea and test its impact on *d_N_/d_S_* which is interpreted as a measure of the efficiency of purifying selection. Contrary to mammals, where an equivalent relationship is strong, we found no relationship between body size and *d_N_/d_S_* in either termites (Spearman’s *ρ* = *−*0.6, *P* = 0.2417) or non-termite Blattodea (Spearman’s *ρ* = 0.1645, *P* = 0.2439). This result converges with what has been observed in another insect clade where eusociality has evolved, the Hymenoptera, whose *d_N_/d_S_* is explained by colony size (Rubin, 2022) and not affected by body size (Benitiere et al., 2022; Weyna and Romiguier, 2021).

We then test the hypothesis that an increase in social organization within the Blattodea would reduce the effectiveness of natural selection by reducing the number of reproductive individuals. To do this, we assigned each of the 66 species studied to one of the social categories defined by Michener (1969).

- Solitary = no social behaviour, individuals live separately from each other.
- Gregarious = individuals live together but do not co-operate or care for young.
- Subsocial = individuals take care of their young for a certain period of time.
- Communal = individuals of the same generation cooperate in nest building but not in the care of young.
- Quasisocial = individuals from the same generation use the same nest and cooperate in the caring for the young.
- Semisocial = individuals from the same generation cooperate in caring for young and sharing reproductive tasks.
- Eusocial = individuals from several generations cooperate in caring for the young and where reproductive tasks are shared.

Throughout the explored dataset, we found 24 solitary species, 31 gregarious, 3 subsocial and 6 eusocial (social organization was not assigned for species of the genus *Tanzaniophasma* and for *Tryonicus parvus*). Statistical analyses highlighted a strong effect of the social system (ANOVA test, *P <* 2.66 *×* 10*^−^*^16^; Fig. 2-B; Table 1).

**Table 1.** Tukey multiple comparisons of means for *d_N_/d_S_* ratios across different social systems. The table displays the pairwise comparisons between each social system. The *difference* column shows the difference in mean *d_N_/d_S_* ratios between the two groups being compared. The *lower* and *upper* columns represent the lower and upper bounds of the 95% confidence interval for these mean differences, respectively. The *adjusted p-values* indicate the significance of these differences after adjusting for multiple comparisons. Significant p-values (p < 0.05) are highlighted in bold.

| Comparison | Difference in Means | Lower Bound | Upper Bound | Adjusted $P$ |
| --- | --- | --- | --- | --- |
| Subsocial - Gregarious | 0.0011 | -0.0110 | 0.0133 | 0.9948 |
| Solitary - Gregarious | -0.0028 | -0.0083 | 0.0027 | 0.5327 |
| Eusocial - Gregarious | 0.0386 | 0.0296 | 0.0475 | **<0.0001** |
| Eusocial - Subsocial | 0.0374 | 0.0232 | 0.0517 | **<0.0001** |
| Eusocial - Solitary | 0.0414 | 0.0322 | 0.0506 | **<0.0001** |
| Solitary - Subsocial | -0.0039 | -0.0163 | 0.0084 | 0.8338 |

The results of a Tukey-HSD test (Table 1) suggest that the eusocial social system is associated with a higher *d_N_/d_S_* than the other groups (gregarious, subsocial, and solitary). The other comparisons (subsocial *versus* gregarious, solitary *versus* gregarious, and solitary *versus* subsocial) did not show statistically significant differences, indicating that *d_N_/d_S_* is only increased for eusocial species. At this phylogenetic scale, our analysis is potentially influenced by a pronounced phylogenetic bias, given the fact that only termites are classified as eusocial in Blattodea. We therefore note:

1. *d_N_/d_S_* is a rate along a branch and is estimated in a phylogenetic framework, which means that technically two neighbouring branches are independent and should not have similar values due to phylogenetic inertia in the same way than two neighbouring nodes for other statistics (such as %*GC* or body mass).
2. When adding this result with those obtained in a very phylogenetically distant group, Hymenoptera (Weyna and Romiguier, 2021), this is the fourth observation of strongly increased *d_N_/d_S_* associated to the emergence of eusociality.

Our results further support the idea that eusociality is linked to a genome-wide reduction in the efficiency of natural selection. This is evidenced by the RELAX analysis, which indicates that approximately 32% of the 3,166 coding genes studied are strongly consistent with a model of relaxed selection, whereas only about 0.72% of the genes are strongly supported by a model of positive selection (Fig. S1).

Interestingly, we note that the closest relative of termites, *Cryptocercus wrighti*, displays an intermediate *d_N_/d_S_* ratio highter to non-social species and relatively close to eusocial ones (*d_N_/d_S_* = 0.0905 compared to an average of 0.0675 in non-social Blattodea and 0.107 in eusocial ones). The *Cryptocercus* genus is characterized by biparental parental care of the brood (Evangelista et al., 2019), which likely precede the evolution of eusociality where parental care is extended to sibling workers. This is reminiscent of the situation in *Anthophila* bees, where solitary bees features high parental care (nest building and pollen collection for few larvae) and display elevated *d_N_/d_S_*, suggesting low effective population size preceding multiple evolution of eusociality in this group (Weyna and Romiguier, 2021). Demographic inferences using PSMC-like approaches (H Li and Durbin, 2011; Lynch, Haubold, et al., 2020; Terhorst et al., 2017) in non-social species with high *d_N_/d_S_* ratios, such as *Cryptocercus* and solitary bees, as well as in closely related eusocial groups, could provide a clearer understanding of the demographic context in which eusociality evolved. This would be particularly valuable for pinpointing the timing of *Ne* variations associated with the emergence of eusociality. This approach has already been applied to social spiders of the genus *Stegodyphus*, where it revealed a reduction in *Ne* associated with increasing social complexity (Ma et al., 2024). Moreover, it could help determine whether the reduction in *Ne* preceded the emergence of eusociality in bees (from solitary species) and termites (from *Cryptocercus*), or if it resulted from the evolution toward more complex social structures.

These results reinforce the idea that high parental care and low effective population size might facilitate evolution towards eusociality. As eusociality is a reproductive strategy characterized by care of offspring shared by many siblings, high parental care is likely to be a first pre-requisite to its evolution. High parental care has also been identified as the best ecological predictor of species with low effective population sizes (Romiguier, Lourenco, et al., 2014). As low effective population size increases the probability of sharing common gene ancestry (Wang et al., 2016), it is expected to increase genetic relatedness among individuals and thus amplify the benefits of kin selection (Epplen et al., 1999; Tabadkani et al., 2012). In this regard, the evolution of eusociality in termites might have been facilitated by the combination of high parental care and reduced effective population size. Whether these are universal pre-requisites for eusociality emergence is still open to question: while low effective population size seems to precede sociality in termite and bees, the same pattern has not been observed in ants and social wasps (Weyna and Romiguier, 2021). This can either mean that eusociality can emerge without initial drop of effective population size, or just that we did not detect it because of a lack of genomic data for closely related non-social species for ants and social wasps.

To further understand the influence of sociality on the effectiveness of natural selection in termites, we analyzed an additional dataset derived from transcriptome sequencing (Bucek et al., 2019). This dataset encompasses 68 termite species as well as 7 dictyopteran outgroups, allowing us to test whether the diversity in levels of eusociality among termites correlates with *d_N_/d_S_* variations.

### Contrasting *d_N_/d_S_* ratios among eusocial Blattodea (Isoptera)

In the second part of our study, we investigated the evolution of the *d_N_/d_S_* ratio within Isoptera using transcriptome data from Bucek et al. Similarly to the approach adopted for Blattodea, we computed the median genomic *d_N_/d_S_* for all branches of the Isopteran phylogenetic tree. The reported *d_N_/d_S_* per species corresponds to their respective terminal branch value (Fig. 3).

**Figure 3.**
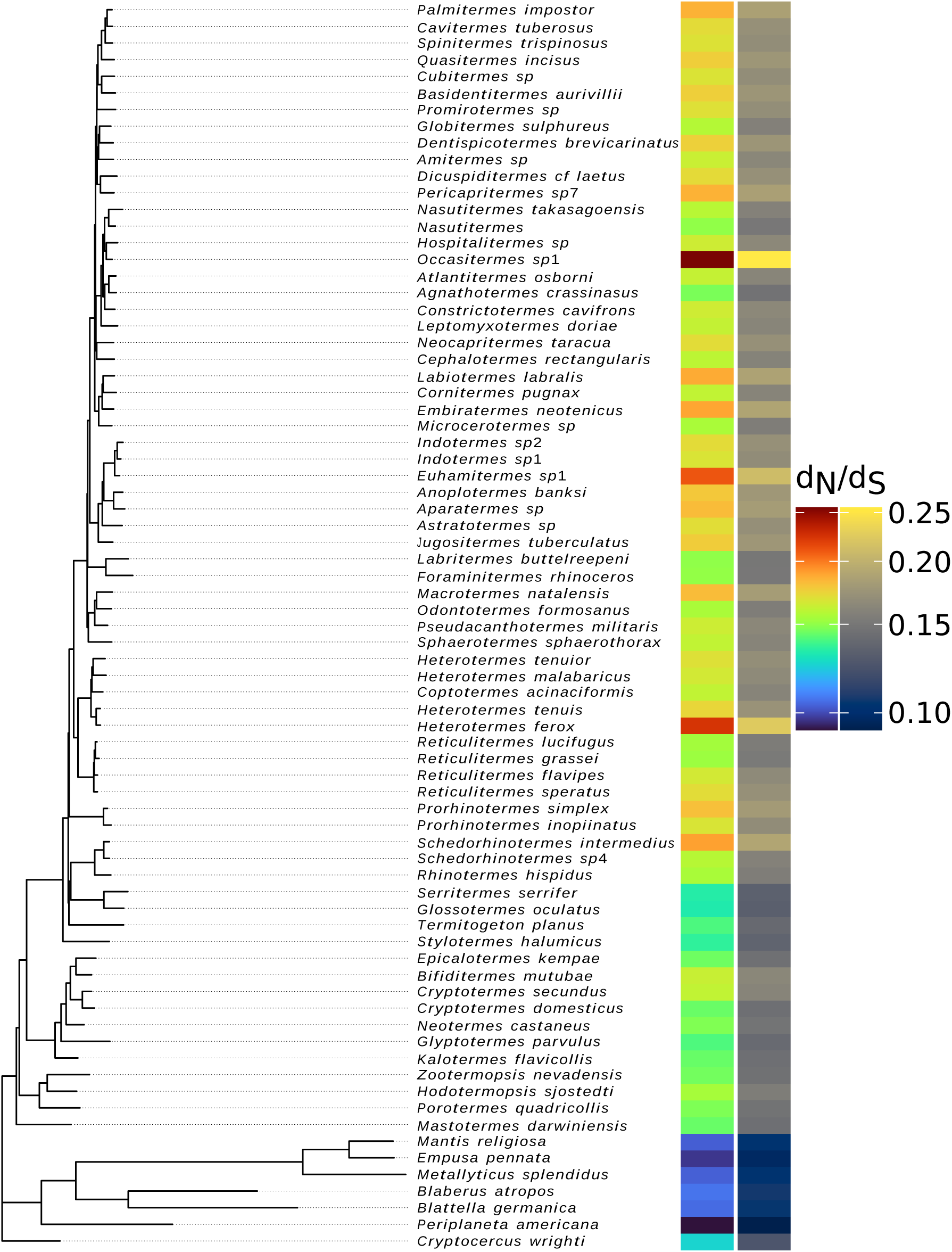
Distribution of the *d_N_/d_S_* ratio in Isoptera (and outgroup species). The phylogenetic tree employed in this study was produced by Bucek et al. (2019), constructed from sequenced transcriptomes. Utilizing the identical coding genes, we estimated the *d_N_/d_S_* ratios along each phylogenetic branch by mapnh. In this study, we exclusively report the *d_N_/d_S_* values corresponding to the terminal branches, represented here by a gradient of colours between 0.0925 (for the outgroup *Periplaneta americana*) and 0.256 (*Occasitermes*). Two color palettes are used here to include the diversity of color perception.

The transcriptomes were available for a total of 75 species. Of these, 68 belong to the Isoptera, exhibiting a *d_N_/d_S_* variation range from 0.132 (*Glossotermes oculatus*) to 0.256 (*Occasitermes*). The remaining seven species constitute the outgroup, showing a *d_N_/d_S_* variation range from 0.0925 (*Periplaneta americana*) to 0.125 (*Cryptocercus wrighti*). This second dataset thus corroborates the overall trend we previously identified using the first dataset (Evangelista et al., 2019), also indicating an increased *d_N_/d_S_* in Isoptera, with values that do not overlap with the range found in non-eusocial Blattodea.

We then tried to explain the large variations of *d_N_/d_S_* across termites. More particularly, we hypothesized that the effect of eusociality on *d_N_/d_S_* could be modulated by the level of social complexity (Table S2). As in ants (Romiguier, Borowiec, et al., 2022), termite societies can be more or less complex, with some species featuring extreme forms of eusociality with highly specialized division of labour, large and long-lived colonies. To test whether social complexity increase the effect of eusociality on effective population size (*Ne*), we tested the effect on *d_N_/d_S_* of two social traits recognized as two good proxies for social complexity in the literature: nesting strategies and developmental pathways (Mizumoto and Bourguignon, 2021). Nesting strategies are typically divided in three categories following Abe (1987): OP (One Piece), MP (Multiple-Piece), and SP (Separated-Piece). In the One-Piece strategy, termites inhabit a single piece of wood that serves as both their nest and food source, resulting in typically small, short-lived colonies. Multiple-Piece nests involve colonies that occupy and connect several pieces of wood via excavated tunnels or constructed shelter tubes, allowing safe travel between different nest sites. In contrast, Separated-Piece nests are physically detached from the food sources, often featuring complex internal structures like chambers and corridors, and are commonly found in subterranean, mound, or arboreal forms (Mizumoto and Bourguignon, 2020). MP and SP colonies are typically larger than OP, but social complexity is at its highest in Separated Piece strategies, with large nests distinct from food source, meaning that they do not reduce in size over time and can growth in size over extended period of time. We conducted a Mann-Whitney U test between each pair of nesting strategies, and always found significant differences except for the MP *versus* SP comparison (*W* =211; adjusted *P* =1; Table S3). Although termites, collectively, exhibit a higher *d_N_/d_S_* ratio compared to non-social Blattodea species, the variations within termites are strongly associated with the complexity of their eusocial structures. Larger colonies, characterized by multiple pieces (MP) and separated pieces (SP) strategies, display higher *d_N_/d_S_* values than the smaller, single-piece (OP) colonies (Fig. 4-A). Within the same termite family, different species tend to exhibit identical nesting strategies, complicating the attribution of a direct correlation between nesting strategy and *d_N_/d_S_*. Nonetheless, a minimum level of variation in nesting strategies is at least observed within two families: Archotermopsidae, with two OP species and one MP species, and Rhinotermitidae B, comprising two OP species and three MP species. Despite the limited number of data points, we observed that within these families, the *d_N_/d_S_* ratio is consistently higher in MP species compared to OP (Fig. S2). This preponderance of species with less complex sociality in the Archotermopsidae could explain why it is the termite family with the lowest *d_N_/d_S_* (median=0.149) over the 11 surveyed families (Table S4).

**Figure 4.**
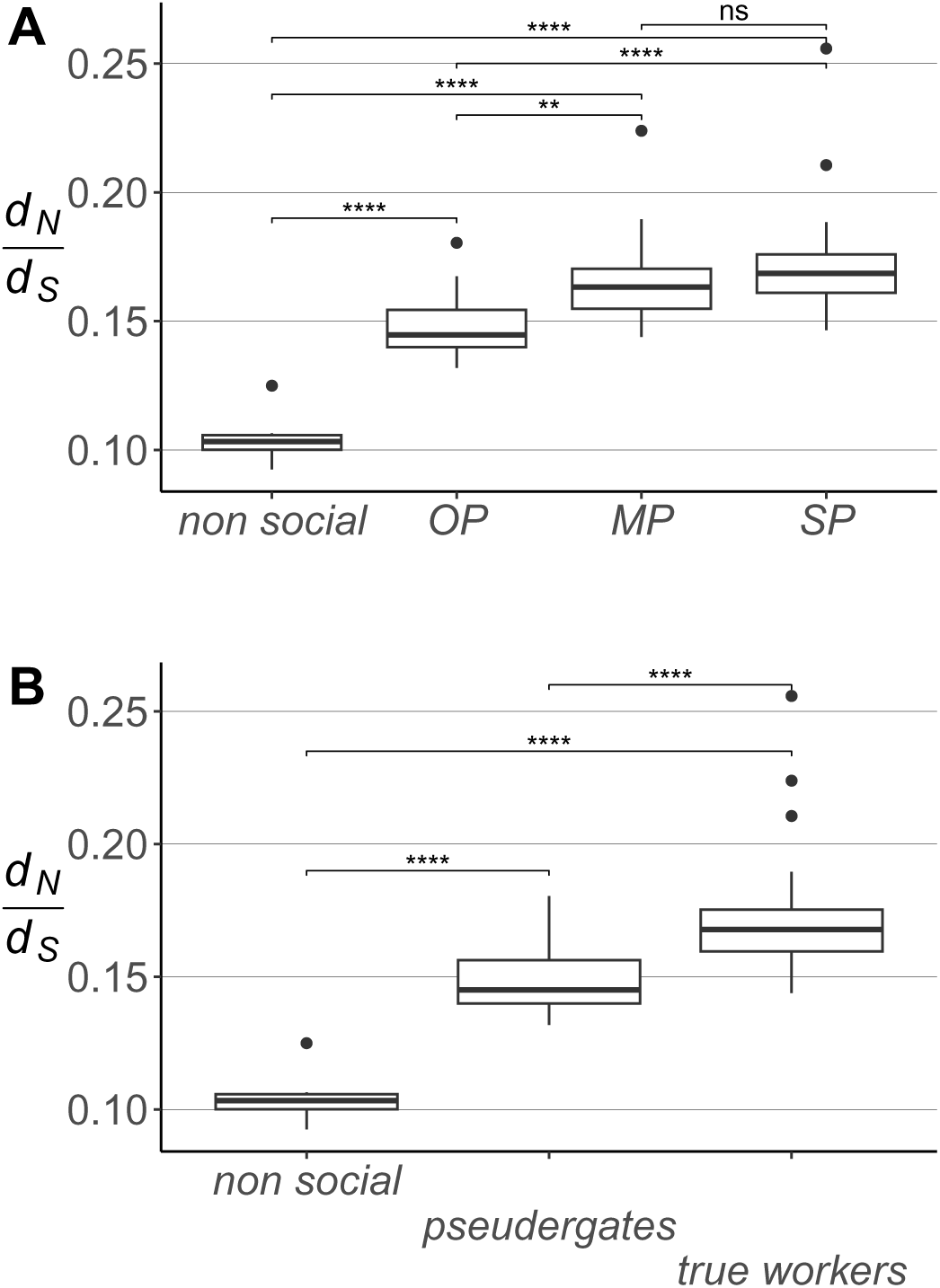
Relationship between social complexity in termites and *d_N_/d_S_* ratio. A. Social complexity, as approximated by nesting strategy, following Mizumoto and Bourguignon (2021). Non-social species are represented solely by the outgroup. OP denotes One-Piece nesters, MP signifies Multiple-Pieces nesters, and SP refers to Separate-Pieces nesters. B. Social complexity, as approximated by the working castes, following Mizumoto and Bourguignon (2021). Pseudergates are workers with the plasticity to transition between castes, including reproductive roles, whereas true workers lack this ability to change castes. Bonferroni-adjusted statistical significance is denoted as follows: *ns* (*P >* 0.05), * (*P ≤* 0.05), ** (*P ≤* 0.01), *** (*P ≤* 0.001), **** (*P ≤* 0.0001).

This intricate social structure likely leads to higher *d_N_/d_S_*values due to the formation of larger colonies, which in turn reduces the number of breeding individuals, significantly affecting *Ne* and thus natural selection efficiency. Such association between advanced sociality and high *d_N_/d_S_* ratio has been retrieved in two empirical studies (Romiguier, Gayral, et al., 2014; Rubin, 2022) but not in a recent study in ants (Barkdull and Moreau, 2023). In contrast, colonies that inhabit a single piece of wood, referred to as OP (One Piece strategy; see Fig. 4-A), are typically smaller. Consequently, these are anticipated to exhibit lower *d_N_/d_S_* values, stemming from decreased reproductive variance among females.

Mizumoto and Bourguignon (2021) suggests an alternative method for gauging the complexity of eusociality in termites by classifying species based on the developmental pathways of their worker caste, which are either pseudergates or true workers. Pseudergates are developmentally immature reproductive individuals, and can perform the colony’s labor tasks but also differentiate later into either soldiers or reproductives. In contrast, true workers lack such developmental plasticity and are highly specialized to the colony’s tasks. This specialization of true workers, as opposed to pseudergates, results in a higher division of labor, thereby indicating increased social complexity. Both species possessing pseudergates or true workers exhibit higher *d_N_/d_S_* ratios compared to non-social species (Mann-Whitney *U* test: *W* =0 in both comparisons; Adjusted *P* -value of 6.26 *×* 10*^−^*^4^ and 6.29 *×* 10*^−^*^5^ respectively; Fig. 4-B). This reinforces the idea that effective population size is not only affected by eusociality, but also at a more finer scale by the different levels of social complexity that affect effective population size. One potential effect increasing further *d_N_/d_S_* in “true workers” species is that worker being sterile, worker-specific genes can not be directly selected but only *via* kin selection. Such an indirect selection is predicted to be reduced compared to direct selection, an idea that has been supported by previous work in the pharaoh ant (Warner et al., 2017). To better quantify the effect of indirect selection in termites and disentangle it from the effect of *Ne*, an interesting perspective would be to replicate our study on queen-biased genes *vs.* worker-biased genes.

An effect of worker reproduction has also been tested in a recent study in ants but led to different results (Barkdull and Moreau, 2023). We suggest that this difference between ant and termites can be explained by differences in turnover between these two groups. In termites, species with true workers are highly represented in one clade which suggests that this trait is highly stable throughout their evolution. In contrast, worker polymorphism in ant species may be less stable as suggested by Barkdull and Moreau (2023). We therefore propose that worker polymorphism in ants might not be maintained over long enough periods to have as clear effects as in termites on *d_N_/d_S_* ratio.

Overall, our results suggest with two different proxies of eusociality level that there is a molecular cost to social complexity. Such a finding have been first suggested in ants, where a correlation between *d_N_/d_S_* ratios and another proxy of social complexity (queen/worker dimorphism) have been reported (Romiguier, Lourenco, et al., 2014). Taken independently, these results in ants and termites are sensitive to correction for phylogenetic inertia, as species with high social complexity tend to be clustered in few related clades. Taken collectively, both results therefore reinforce the conclusion of the other, as it seems unlikely that the association between high *d_N_/d_S_* and high social complexity is due to chance in both distant taxa. Further reinforcing this conclusion, an association between advanced sociality and a high *d_N_/d_S_* ratio has also been observed in bees (Rubin, 2022), indicating that this pattern is present across all three major eusocial taxa. A similar trend has been identified in social *Synalpheus* shrimps compared to their non-social counterparts (Chak et al., 2021). These findings provide yet another example of the negative impact of eusociality on the efficiency of natural selection.

We therefore note that less efficient purifying selection at low *Ne* does not necessarily translate into less positive selection events. In bees, eusocial lineages appear to exhibit higher *d_N_/d_S_* ratio (likely due to less efficient purifying selection), but also more positive selection events (Shell et al., 2021). In a similar way, a positive selection hotspot is associated to the origin of advanced eusociality in ants (Romiguier, Borowiec, et al., 2022). Such an apparent paradox can be explained because smaller populations are predicted to have a larger proportion of beneficial mutations due to the increased fixation of deleterious mutations in such populations, which in turn creates more opportunities for new beneficial back-mutations (i.e, compensatory mutations; Weissman and Barton, 2012). This theoretical expectation of saturation in adaptive rates over the long term for species with large *Ne* has been empirically demonstrated across various animal groups (Rousselle et al., 2020), including insects, mollusks, annelids, birds, and mammals. In this study, the relationship between effective population size and *ω_A_*, the rate of adaptive amino-acid substitution is indeed positive only in taxa with low *Ne*.

To conclude, our study supports that eusociality is associated to significant decreases in terms of natural selection efficiency, which validates population genetic theory and highlights that eusocial insects undergo molecular evolution that is closer to large vertebrates than other insects. This support the view that social insects should be considered as superorganisms (Boomsma and Gawne, 2018), and that their population size vary depending on their social complexity in the same way than population size vary depending on life history traits for non-social species (Romiguier, Gayral, et al., 2014).

## Data, script, code, and supplementary information availability

Data and scripts are available online from a Zenodo depository: https://doi.org/10.5281/zenodo.11057544

---

**Figure S1.**
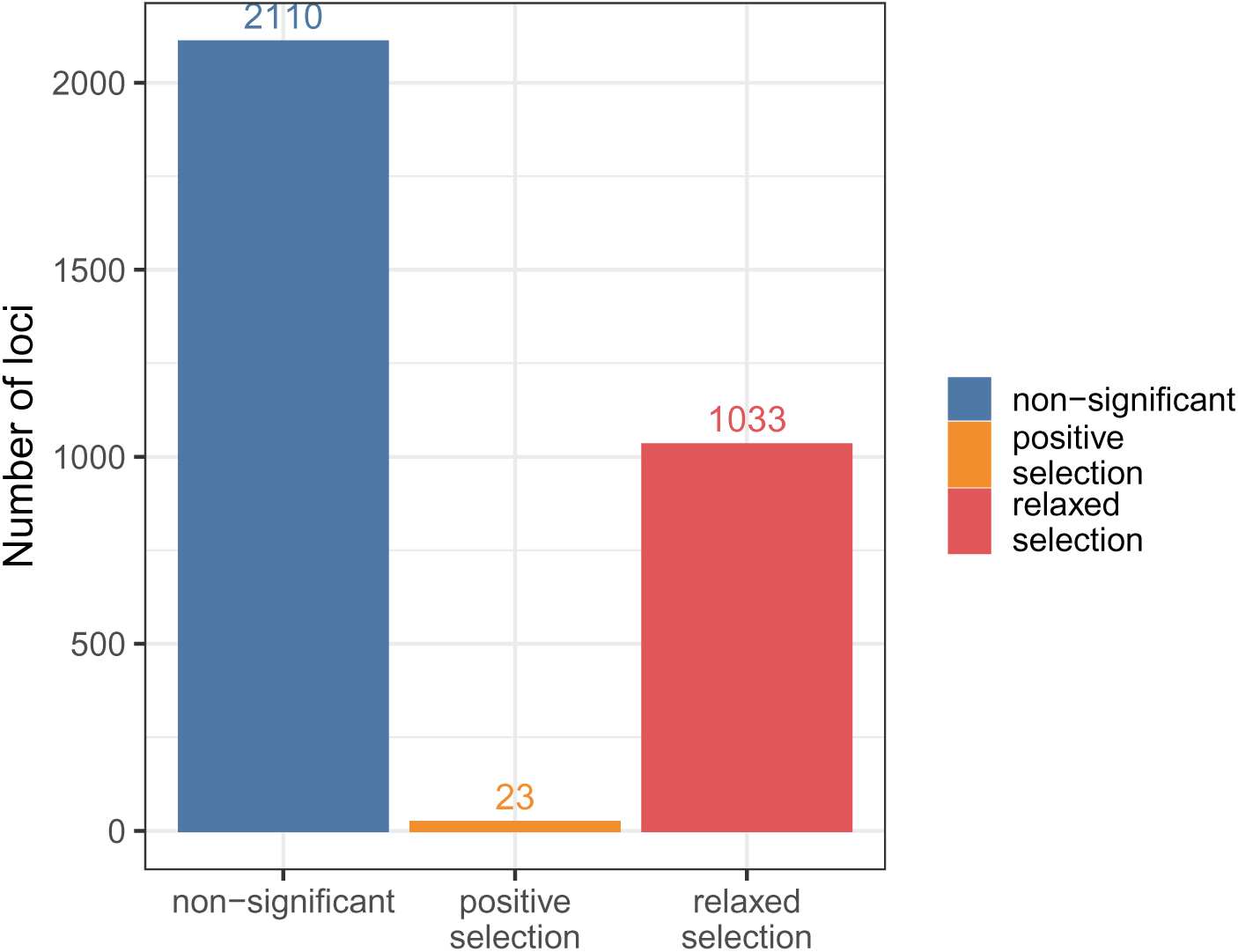
Test for the relaxation of purifying selection (*k <* 1) and positive selection (*k >* 1) against a null model for 3,166 orthologous codon-aligned genes in the Evangelista et al. (2019) dataset. Blue: loci that do not reject the null model (p-value > 0.05 after Bonferroni correction). Orange: loci supporting a positive selection model in termites (*k >* 1, p-value < 0.05 after Bonferroni correction). Red: loci supporting a relaxation of purifying selection in termites (*k <* 1, p-value < 0.05 after Bonferroni correction).

**Figure S2.**
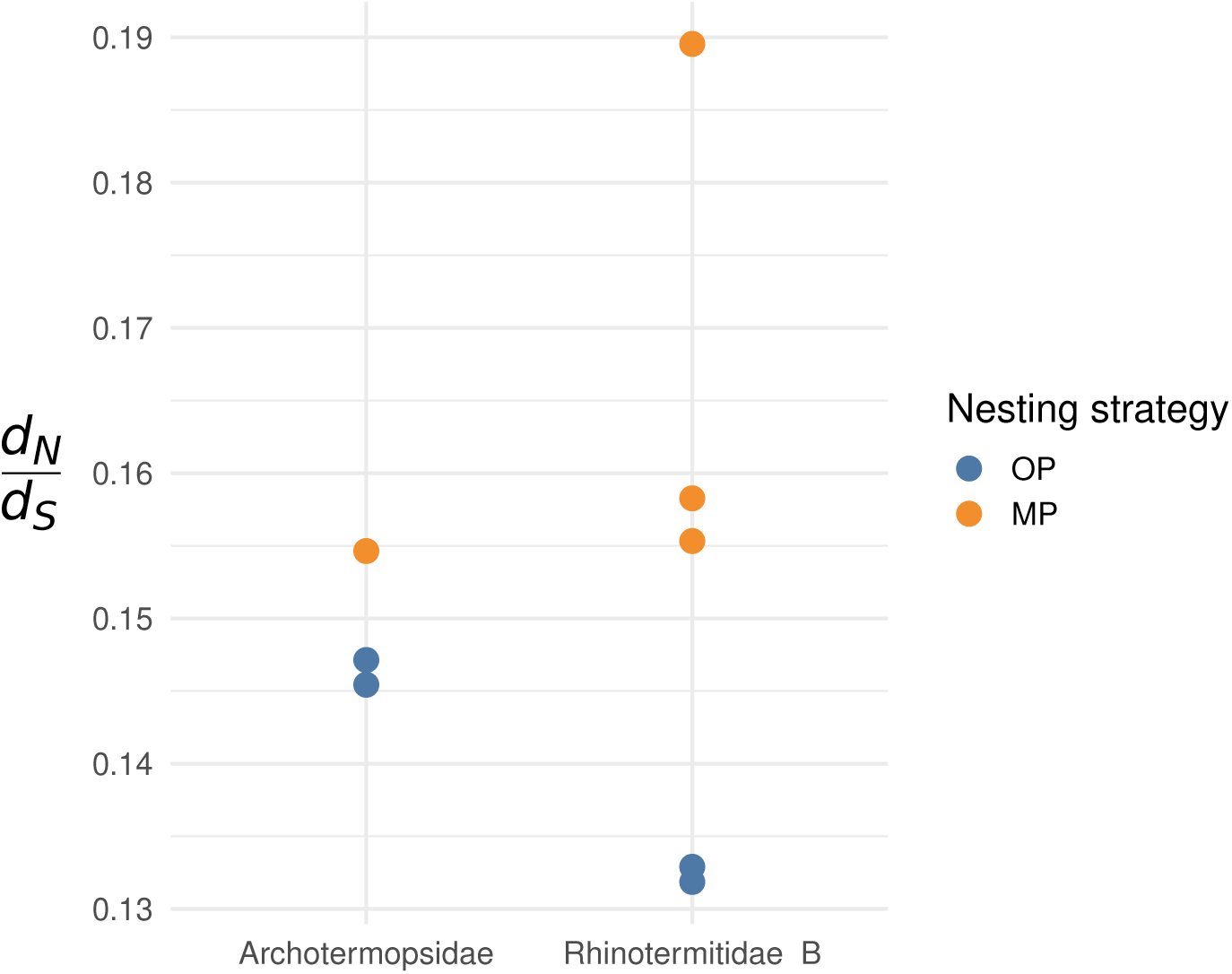
Association between social complexity and *d_N_/d_S_* ratios in Archotermopsidae and Rhinotermitidae B using the nesting strategy as proxy.

**Table S1.**
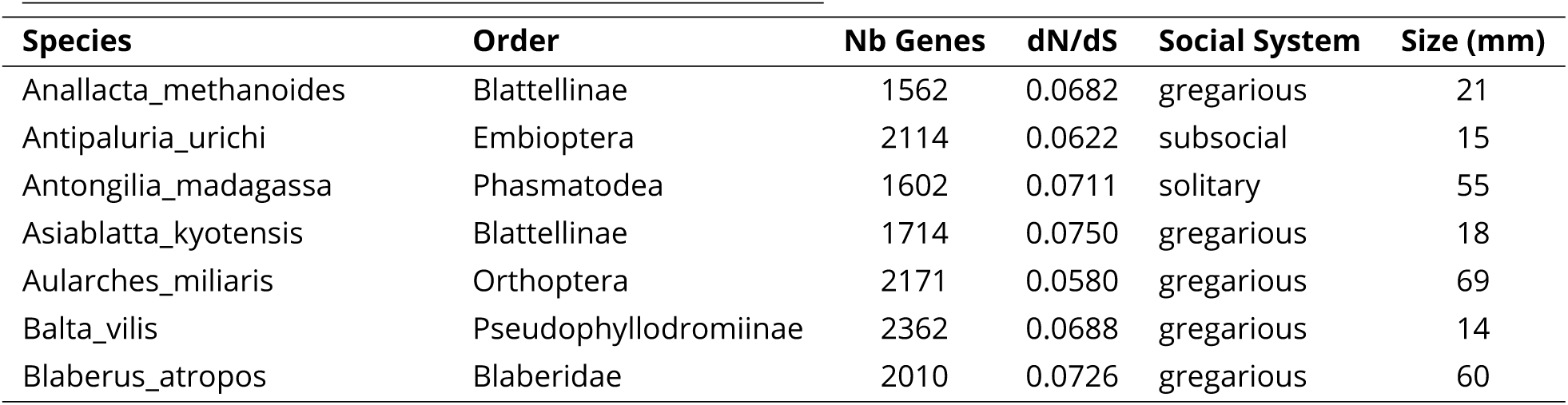
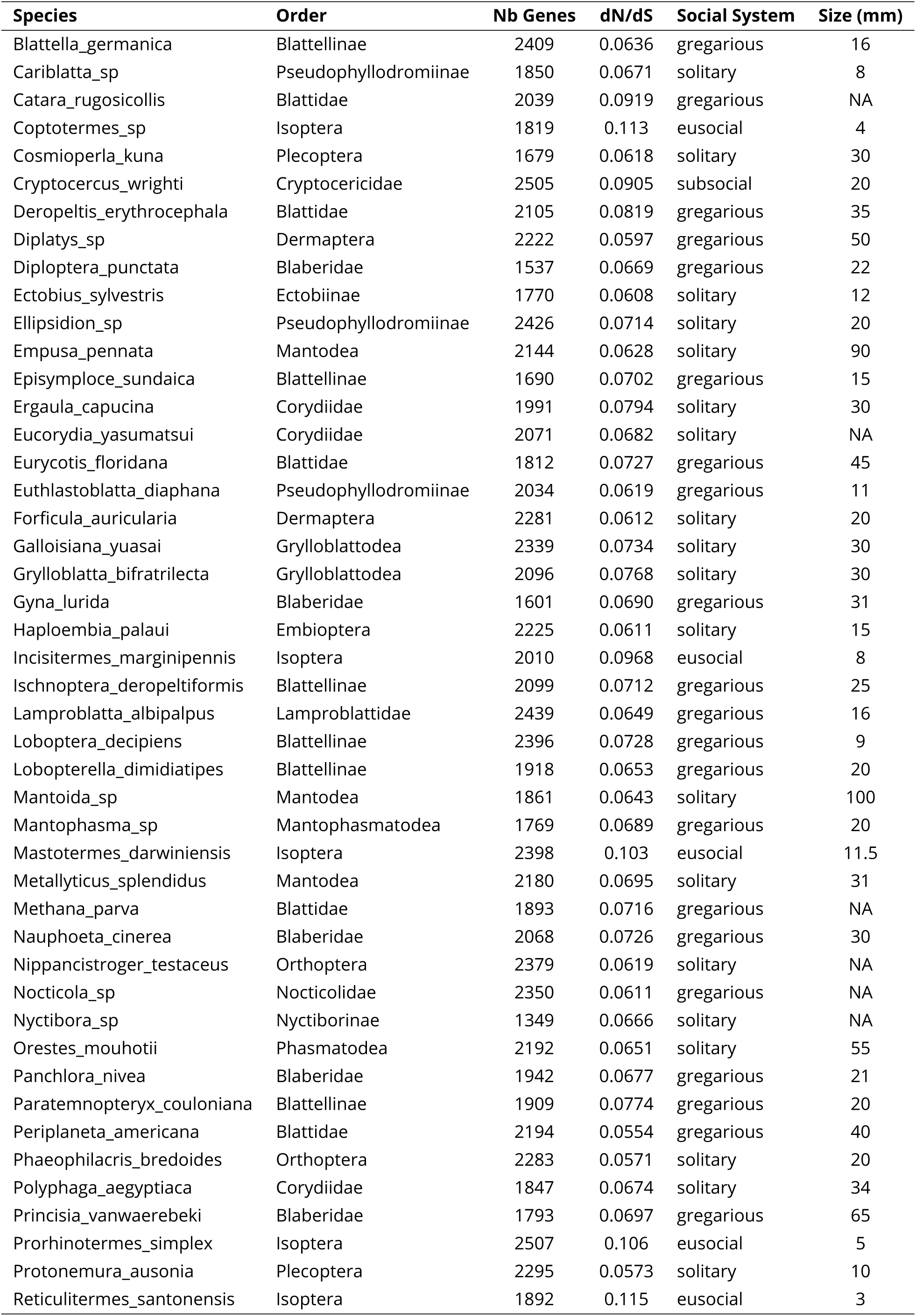
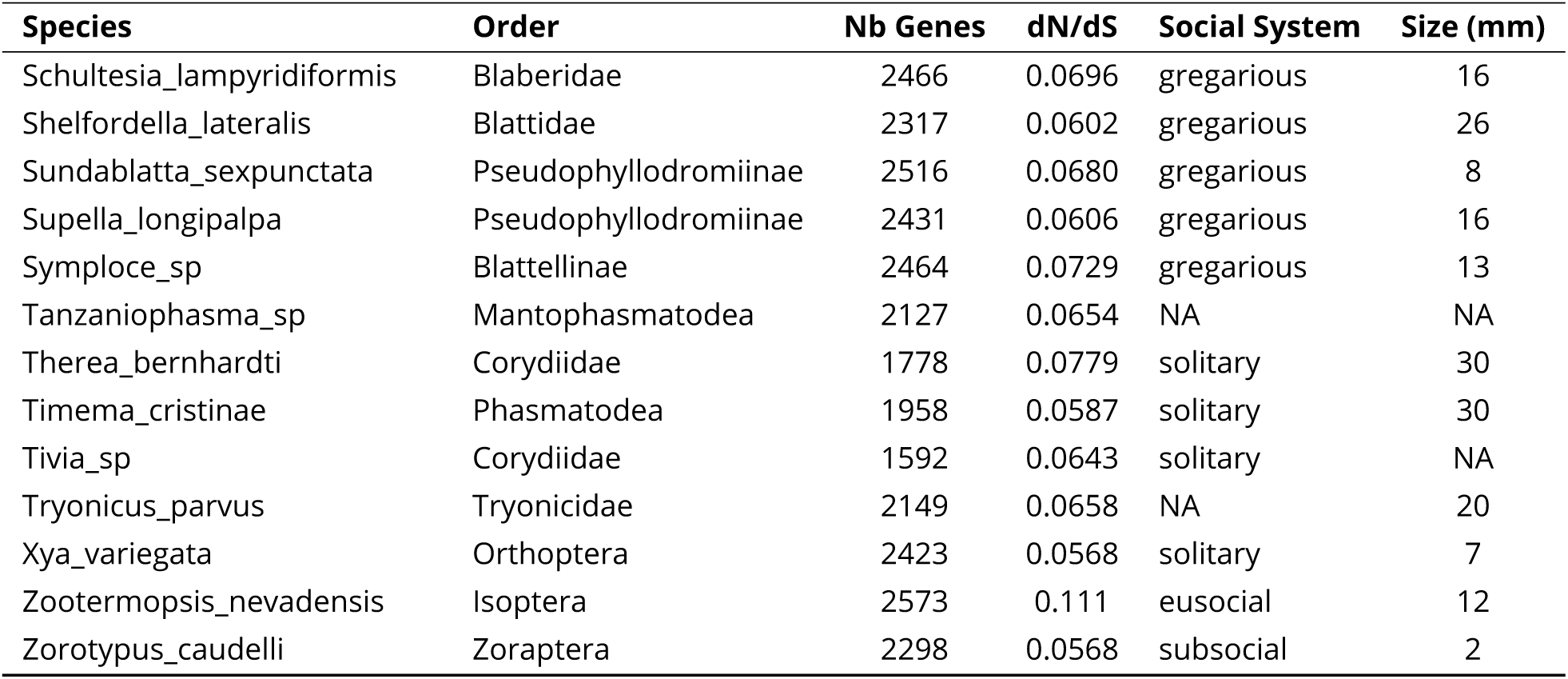
Relationship between *d_N_/d_S_* ratios, social traits and adult body size in the Blattodea dataset (Evangelista et al., 2019)

**Table S2.**
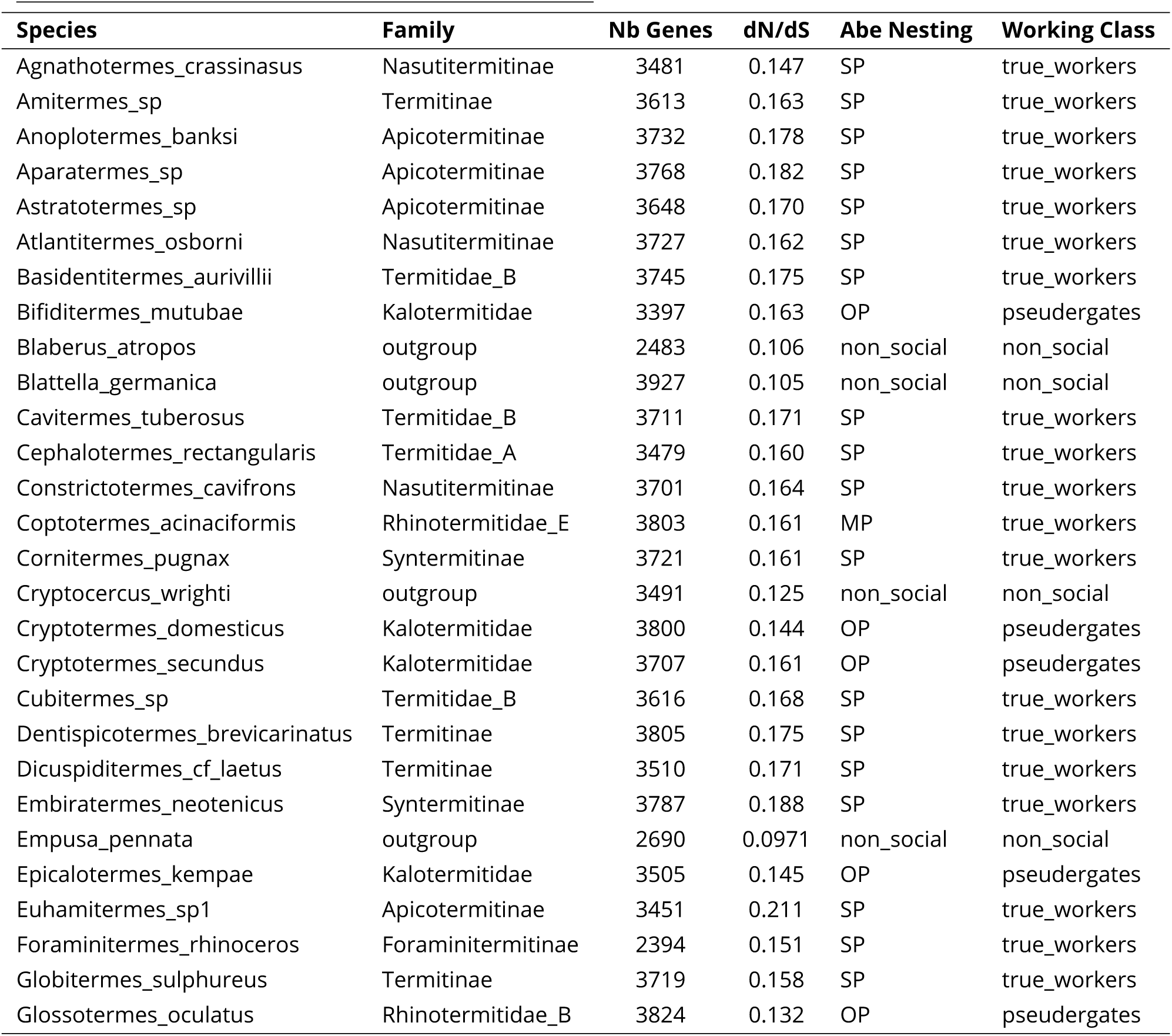
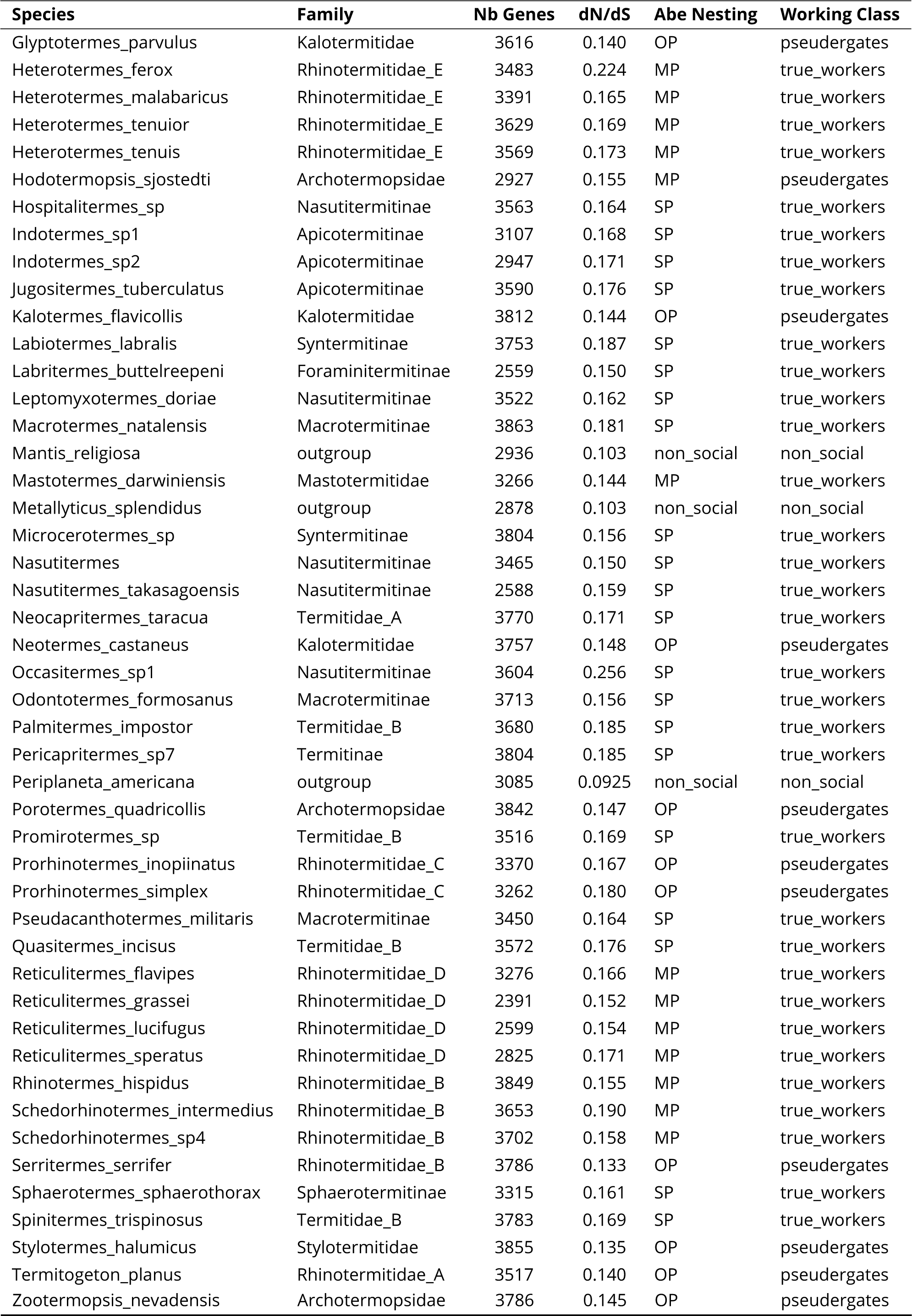
Relationship between *d_N_/d_S_* ratios and social traits in the Isoptera dataset (Bucek et al., 2019)

**Table S3.** Pairwise comparisons of differences in *d_N_/d_S_* between alternative nesting strategies

| Pair | *W* | Adjusted *P*-value |
| --- | --- | --- |
| Non-social vs. OP | 0 | $1.48 \times 10^{-3}$ |
| Non-social vs. MP | 0 | $1.78 \times 10^{-3}$ |
| Non-social vs. SP | 0 | $1.91 \times 10^{-4}$ |
| OP vs. MP | 40 | $2.93 \times 10^{-2}$ |
| OP vs. SP | 74 | $1.53 \times 10^{-4}$ |
| MP vs. SP | 211 | 1.00 |

Adjusted *p*-values from Mann-Whitney *U* tests comparing the $d_N/d_S$ values between different nesting strategies of termite species. Adjustments were made using the Bonferroni method to correct for multiple comparisons.

**Table S4.** *d_N_/d_S_* values for termite families

| Family | mean $d_N/d_S$ | min $d_N/d_S$ | max $d_N/d_S$ |
| --- | --- | --- | --- |
| Apicotermitinae | 0.179 | 0.168 | 0.211 |
| Archotermopsidae | 0.149 | 0.145 | 0.155 |
| Kalotermitidae | 0.149 | 0.140 | 0.163 |
| Macrotermitinae | 0.167 | 0.156 | 0.181 |
| Nasutitermitinae | 0.170 | 0.147 | 0.256 |
| Rhinotermitidae_B | 0.154 | 0.132 | 0.190 |
| Rhinotermitidae_D | 0.161 | 0.152 | 0.171 |
| Rhinotermitidae_E | 0.178 | 0.161 | 0.224 |
| Syntermitinae | 0.173 | 0.156 | 0.188 |
| Termitidae_B | 0.173 | 0.168 | 0.185 |
| Termitinae | 0.170 | 0.158 | 0.185 |

